# A point mutation in Chl biosynthesis gene Mg-chelatase I subunit influences on leaf color and metabolism in strawberry

**DOI:** 10.1101/2022.06.30.498292

**Authors:** Yang-Yang Ma, Jian-Cheng Shi, Dan-Juan Wang, Xia Liang, Feng Wei, Chun-mei Gong, Li-juan Qiu, Ying-Qiang Wen, Jia-Yue Feng

## Abstract

Magnesium chelatase catalysis the insertion of magnesium into protoporphyrin IX is a vital step in chlorophyll biogenesis. It consists of three subunits, CHLI, CHLD and CHLH. The CHLI subunit is an ATPase and hydrolysis ATP in the catalysis. However, its key point on influencing flavonoid biosynthesis and chlorophyll accumulation under different light density was still unknown. In this study, we identified an N-Ethyl-N-nitrosourea (ENU) mutant, p240 from strawberry Fragaria pentaphylla that has yellow-green leaves and lower chlorophyll level. We verified the mutation occurs in the 186th amino acid of CHLI subunit, which is conserved in most photosynthetic organism. Mutants generated from RNAi and CRISPR/Cas9 gene editing confirmed this phenotype. In addition, we found that FpCHLI was localized in chloroplast and its subcellular location have not been changed by mutation. Further study showed that the interaction between FpCHLI and FpCHLD were not affected by mutagenesis. In contrast, all types of mutants showed reduced ATPase and magnesium chelatase activity indicating mutagenesis decreased enzymatic activities. Furthermore, mutagenesis suppressed the biosynthesis of Mg-proto IX. Metabolites analysis of gene knock-out mutant and WT revealed that CHLI may help to keep stabilizing the flavonoid level in leaves. Furthermore, both p240 and chli mutants are light sensitive, which has yellow leaves under high light but pale green leaves under poor light. The photosynthesis ability of mutants was also increased under shade. Moreover, stomatal apertures of mutants were wider than WT. Taken together, these results suggest that CHLI plays an important role in both leaf coloration and metabolism.

## Introduction

Chlorophylls are essential for oxygen photosynthetic organisms and play vital roles in photosynthesis. They can absorb light energy, feed electrons into the photosynthetic electron transfer chain, and drive charge separation reactions in reaction centers (Chen, 2014). Chlorophylls and heme are all tetrapyrrole molecules originated from a common compound, glutamyl-tRNA^glu^. The branch point of biosynthetic pathway is protoporphyrin IX, which leads to ‘iron-branch’ to formation heme in the presence of Fe^2+^ and leads to ‘Mg-branch’ to formation chlorophylls when inserted Mg^2+^(Brzezowski et al.,2015). The insertion of Mg^2+^ into protoporphyrin IX is the first committed step of Chl biosynthesis (Masuda et al., 2008).

Mg chelatase is a key regulator of chlorophyll biosynthesis which catalyzes the insertion of Mg^2+^ into protoporphyrin IX. Mg chelatase belongs to AAA^+^ chelatases and consists of three subunits, I, D and H. These subunits are conserved in plants, algae, *Rhodobacter capsulatus* and other bacteria. In bacteria, these subunits are named BchH, BchD and BchI, while in algae and plants are named CHLH, CHLD and CHLI (Chew et al., 2007; Willows et al., 1996). The ChlH subunit is a porphyrin-binding subunit of Mg chelase (Willows et al., 1996; Qian et al., 2012). Recently, glutamate (E660) in ChlH subunit was reported to be the key catalytic residue for magnesium insertion, into proximity with the porphyrin (Adams et al., 2020). The I and D subunit are members of the AAA+ (ATPases associated with various cellular activities) superfamily, but only I subunit has an active ATPase activity. In the presence of Mg^2+^ and ATP, I subunit forms a complex with D subunit. The ChlID complex interact with ChlH chelation subunit to form an active chelatase enzyme (Farmer et al., 2019). A proline-rich and acidic amino acid domain were identified in ChlD to stabilize the I-D complex (Lundqvist et al., 2010).

The ChlI (BchI) subunit, which belongs to AAA+ ATPase, is responsible for ATP hydrolysis (Lundqvist et al., 2010). The AAA+ ATPase module contains two major domains. The N-terminal subdomains include highly conserved amino acid residues, Walker A and Walker B motifs. The Walker A motif is considered to stablise ATP binding while Walker B matches up with an ATP-bound magnesiumion to promote ATP hydrolysis (Jessop et al., 2021). The C-terminal is a helical bundle domain (Fodje et al., 2001). Analysis of *Rhodobacter capsulatus* BchI and BchD demonstrated that two subunits form a double-ring structure in the presence of ATP, and bind to ChlH with the participation of Mg^2+^ (Fodje et al., 2001).

Several mutants of *CHLI* genes have been reported in higher plants. However, the phenotypes were various according to different mutagenesis position. In *Arabidopsis*, there are two homologous genes *CHLI1* and *CHLI2*, *cs* is a T-DNA insertion mutant in the 3’–terminal end of *CHLI*, resulting a pale-green phenotype (Koncz et al., 1990). *cs215* is a point mutation mutant in CHLI1 which converted a Thr at residue 195 to an IIe (T195I), and the homozygous *cs215* is albino (Huang and Li, 2009). In barley, semidominant mutants *Chlorina-125, -157, and -161* cause single amino acid substitutions at position of L91F, D187N, and R269K, respectively, all these three mutants had yellow, chlorophyll-lacking phenotypes, further study showed they were deficient in ATP hydrolysis and magnesium chelatase activity (Hansson et al; 1999 and 2002). Subsequently, it is proved that ATPase activity of magnesium chelatase subunit I was required to maintain subunit D in vivo in barley (Lake et al., 2004). Rice *Chlorina-9* mutant which indicates a yellowish-green phenotype has a missense mutation at the third exon (R307C) in CHLI, resulting reduced activity of Mg-chellatase (Zhang et al., 2006). A single-base mutation (G177R) in *OsCHLI* was selected as *ell* mutant, which displayed a yellow leaf in young seedlings and became lethal after three-leaf stage. In the mutant, OsCHLI could not interact with the CHLD protein, which disrupted the synthesis of protoporphyrin IX (Zhang et al., 2015).

Semi-dominant *Oil yellow1* (*Oy1-N1989*) mutants of maize exhibits a yellow leaf phenotype and lacks Mg-chelatase activity, and the substitution occurs at position 176, a leucine residue converted by phenylalanine (Sawers et al., 2006). In cucumber, the golden-leaf mutant *C528*, which has an amino acid mutation (G296R) in *CsChlI*, results in reduced magnesium chelation activity and chlorophyll levels (Gao et al.,2016). In wheat, a substitution mutant (D221N) *chli* was identified and exhibited a pale-green leaf, Yeast two-hybrid assay revealed Tachli-7A cannot interact with itself or TaCHLI-7A (Wang et al., 2020). In conclusion, most mutants of CHLI are point mutations, phenotypes are various from pale-green, yellowish-green, yellow to albino. The difference phenotype may be caused by the mutated position in CHLI. In strawberry, there has been none report on malfunctional CHLI mutant and the key position in CHLI still remains unknown.

Chlorophyll, carotenoid and flavonoids are main metabolites in plant leaves. Many yellow leaf mutants revealed a reduced chlorophyll and carotenoids. However, there were few studies on flavonoids changing in yellow leaf mutants. In plants, flavonoids are widely distributed in flowers, leaves and seeds. These compounds can act as antioxidants, by preventing plant cells from damage of reactive oxygen species (ROS) generated from excessive light or fungal and bacterial diseases (Casanova et al., 2020). Because of their structural differences, flavonoids can be classified into seven subclasses: flavonols, flavones, isoflavones, anthocyanidins, flavanones, flavanols, and chalcones (Shen et al., 2022). Flavonoids in leaves are various, catechins which belongs to a class of flavanols is the major flavonoids in tea plant. ‘Huangjinya’ is an light sensitive albino tea germplasm, it has yellow leaves under intense light while green leaves in low light. Studies on flavonoids on ‘Huangjinya’ revealed that catechins were down-regulated in intense light and increased in moderated shading (Song et al., 2017). In ginkgo leaf extracts, flavonol glycosides (quercetin, kaempferol, isorhamnetin) account for 24% of all metabolites. However, the flavonoids subclasses in strawberry leaves still remains unknown.

Insufficient light will affect plant Chl biosynthesis, while excessive light will produce photoinhibition and inhibit Chl synthesis in plants. Evidence from previous studies indicates that proper reduction of light intensity increases chlorophyll accumulation in tea leaves (Song et al., 2012; Wang et al., 2012; Liu et al., 2018) while shade induced accumulation of chlorophylls in *Camellia sinensis* cv. Shuchazao leaves (Liu et al., 2020). However, there were few research on light-related chlorophyll metabolism. Moreover, low light conditions in winter is an important factor limiting yield and quality of strawberry.

In this study, we report a chlorophyll-deficient mutant *p240* from an ENU mutagenesis population of *Fragaria pentaphylla.* We identified its characteristics of phenotypes and candidate genes. We identified a point mutation of magnesium chelatase *CHLI* gene was responsible for phenotype of mutant *p240*. Subsequently, mutants generated from RNAi and CRISPR/Cas9 gene editing were indicated similar phenotype as *p240*. Furthermore, we show that the single amino acid substitution cannot block the intrinsic interaction between itself or FvCHLI and FvCHLD, but results in significantly reduced ATPase and magnesium chelatase activity, suggesting that single amino acid substitution produces inactive complexes. Metabolites analysis of leaves in gene-editing mutant and WT revealed catechin was the major flavonoid in strawberry leaves and mutant showed a significant lower level on catechin. In addition, *chli* mutants exhibited an increase of low light tolerance, higher photosynthesis ability and stomatal aperture than WT. These results provide insight into CHLI function on strawberry chlorophyll and flavonoid metabolism and also for breeding strawberry cultivars with low light tolerance. Furthermore, our study also provides some evidence for the role of CHLI in chloroplast retrograde signaling

## Results

### The *p240* mutant in *F. pentaphylla* shows reduced Chl Level and impaired chloroplast Development

The *p240* mutant was isolated in M_1_ population using an ENU mutagenesis screen in the *F. pentaphylla* white-fruited accession ‘Pao’er’. As shown in Figure 1a, *p240* develops many stolons with large number of daughter plants, just like its parent line. Because of its difficult to flower, we conducted experiments with daughter plants. The mutant and wild type were planted under two different conditions, in a growth chamber (Fig. 1a) with an alternating light/dark cycle (16h of light at 22℃/8h of darkness at 20℃) and a greenhouse (Fig. 1b). Under both conditions, *p240* mutant plants displayed a yellow-green leaf phenotype. In growth chamber, the contents of Chl a and Chl b in *p240* were all less than a third that of the WT, and the carotenoid (Car) was about half that of WT (Fig. 1c and 1d), suggesting that the yellow-green phenotype of *p240* mutant was stable.

**Fig. 1.**
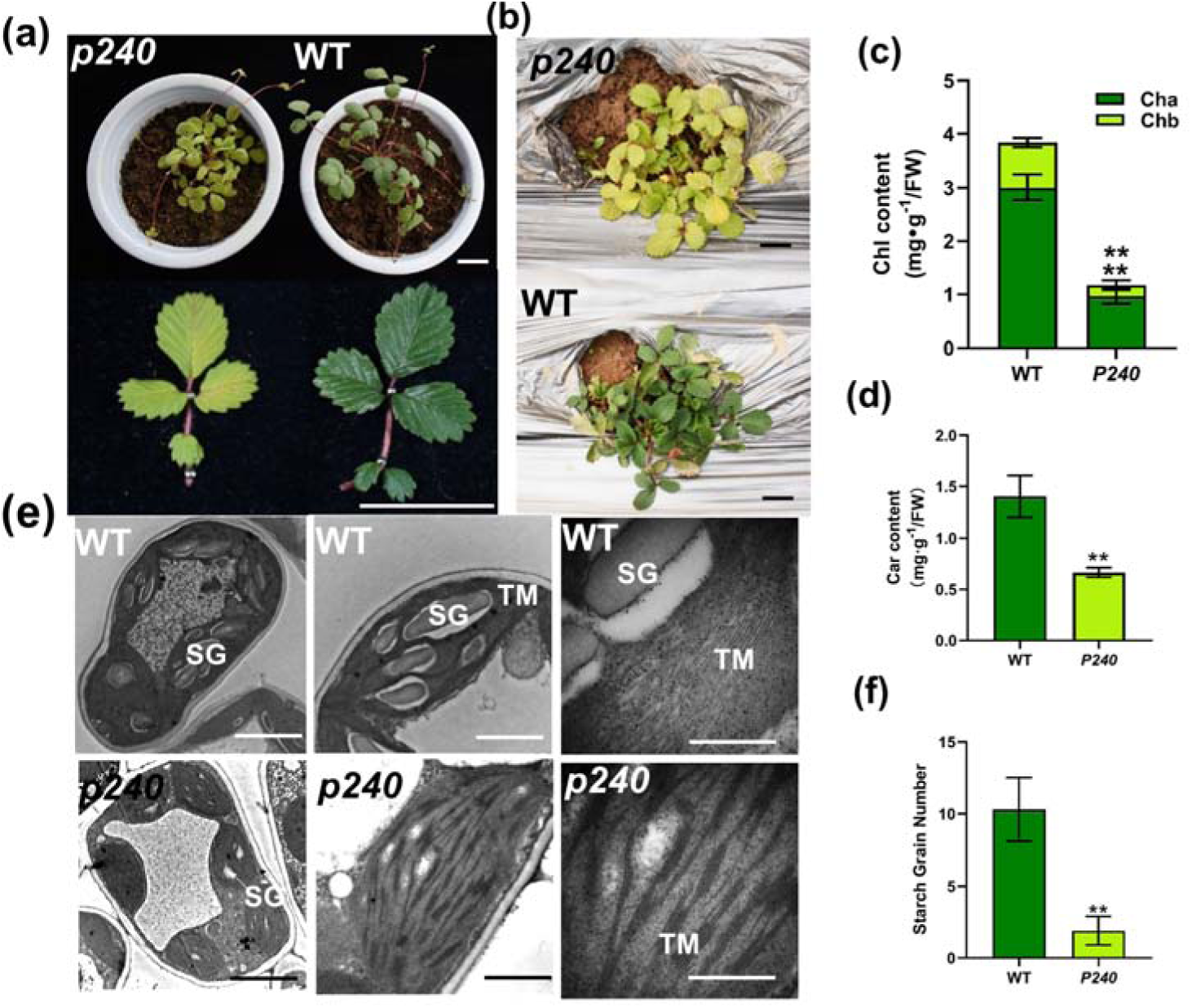
The *Fragaria pentaphylla* ENU mutant *P240* shows yellow-green leaf phenotype. (a) *p240* and WT cultivated in a growth chamber. Bars, 2cm. (b) *p240* and WT cultivated in a greenhouse. Bars, 2cm. (c) Comparison of Chlorophyll contents in WT and *p240*. (d) Carotenoid contents of WT and *p240*. (e) TEM images shows mesophyll cells and chloroplasts of WT and *p240*, from left to right were mesophyll cells, chloroplasts and magnified chloroplasts. Bars, from left to right, 2μm, 1μm, 200nm, respectively. (f) The number of starch grain. SG, starch grain; TM, thylakoid membrane; (c), (d), Data represents the mean±SD of three biological replicates. (f), Data represents the mean±SD of ten biological replicates, all data determined by Student’s t-test. **, *p<0.01*.

Next, whether the yellow-green phenotype was associated with the number and development of chloroplast was investigated. We extracted protoplasts of both *p240* mutant and wild type to calculate the number of chloroplast in one mesophyll cell, and found no significant difference between p240 mutant and wild type (Fig. S1). Transmission electron microscopic (TEM) analysis was conducted to determine development difference of chloroplast. The results showed striking differences in the ultrastructure of chloroplast in *p240* mutant compared to wild type. In the mutant, mesophyll cells had abnormally developed chloroplast with loosely stacked thylakoid membranes (Fig. 1e). In addition, *p240* produced only 20% of wild type starch grain numbers (Fig. 1e and 1f). These results suggested that the gene mutated plays an important role in chloroplast development.

### Isolation and identification of Candidate gene responsing to *p240* phenotype

*F. pentaphylla* prefers growing on high attitude area, and is difficult to flower in flat area. When we screened *p240* and cultivated on greenhouse and growth chamber, we tried various methods to make it flower but we only obtained three M_2_ plants. So it is difficult to perform forward genetics to map gene responsible for *p240* phenotype. Hence we first carried out RNA-Seq analysis to find molecular mechanism of leaf pigments between *p240* and wild type. cDNA libraries were constructed using mature leaves, approximately 40 million reads were obtained for each library and aligned with the *Fragaria vesca* reference genome V4.0 (Edger et al., 2018). The average uniquely mapped efficiency was 79.14% (Table S1). Differentially expressed genes (DEGs) in WT and *p240* were identified. In total, 4557 (DESeq2 padj<=0.05 |log2FoldChange|>=0.0) were detected, where 2333 were upregulated and 2224 were downregulated (Fig.S2a). We performed Go enrichment analysis with these DEGs, and no go term in the ‘cellular component’ was enriched (Fig.S2b). The DEGs also aligned against the KEGG pathway databases. Among 4557 DEGs, 1357(29.8%) were assigned to 114 KEGG pathways. The 19 significant enriched pathways (*p-value* <0.5) were listed (Fig. S2c). Aligned results were also sorted for chromosome coordinates by using picard-tools, the repeated reads were removed, after filtering original data, SNPs were obtained (Fig. S2d).

KEGG enrichment analysis using DEGs revealed associations with porphyrin and chlorophyll metabolism, alanine, aspartate and glutamate metabolism , sulfur metabolism, starch and sucrose metabolism and carbon metabolism, we selected all genes related to these pathways in NCBI database and searched in the SNPs of each samples in RNA-Seq data and compared to find common SNP in *p240*, then we found two genes, ATP-dependent 6-phosphofructokinase 4 (PFK4) and magnesium-chelatase subunit ChlI (CHLI), were mutated (Table S3). To identify the mutagenesis, we cloned the coding region (CDS) of the two genes in *p240* and wild type (WT) and sequenced. The g1028a (E186K) mutation and g3480t (A450S) heterozygous mutation occurred in coding region of *FpChlI* and *FpPFK4*, respectively (Fig. 2a and S3). Further transgenic strawberries of two genes showed *FpPFK4* had no relevance to the phenotype of *p240* and *FpCHLI* may be responsible for yellow-green leaf genotype (Fig. S4, Fig 3 and 4). In addition, three M_2_ *p240* exhibited a green-leaf phenotype (Fig.2b), we also cloned CDS of CHLI in these M_2_ plants, sanger sequencing assays showed that these were all wild-type sequences (Fig. 2a).

**Fig. 2.**
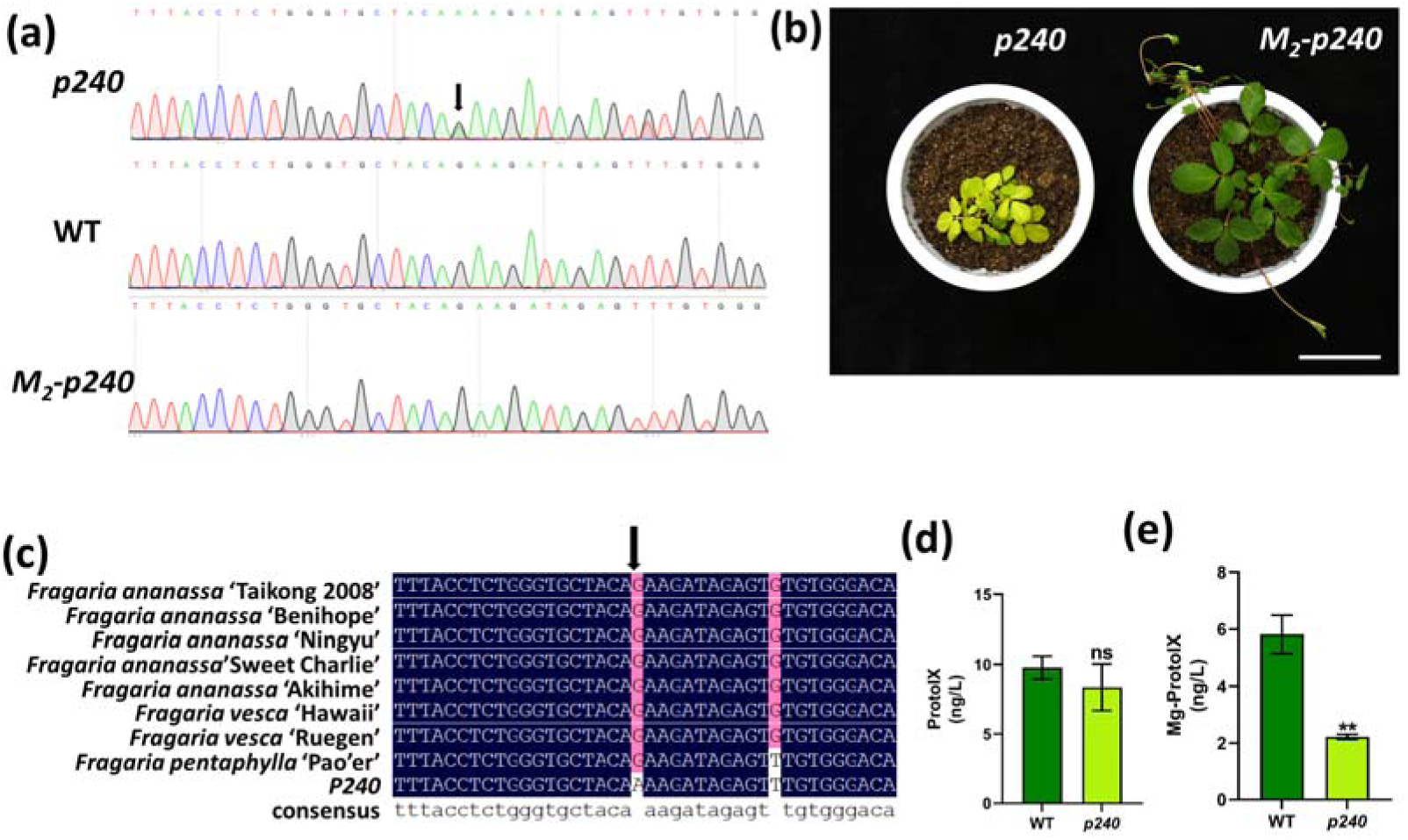
The *p240* mutant in *Fragaria pentaphylla* is caused by a mutation in FpCHLI. (a) Sequencing chromatograms of *CHLI* in *p240*, WT and M_2_-*p240*. (b) Phenotype of M_1_ population of *p240* and M_2_ population of *p240*, bar=5cm. (c) Alignment of CHLI in various strawberries. Blue arrow shows mutated position. (d) Proto IX contents of *p240* and WT. (d) Mg-Proto IX contents of *p240* and WT. Data represents the mean±SD of three biological replicates. all data determined by Student’s t-test. **, *p<0.01*, ns, no significant difference.

**Fig. 3.**
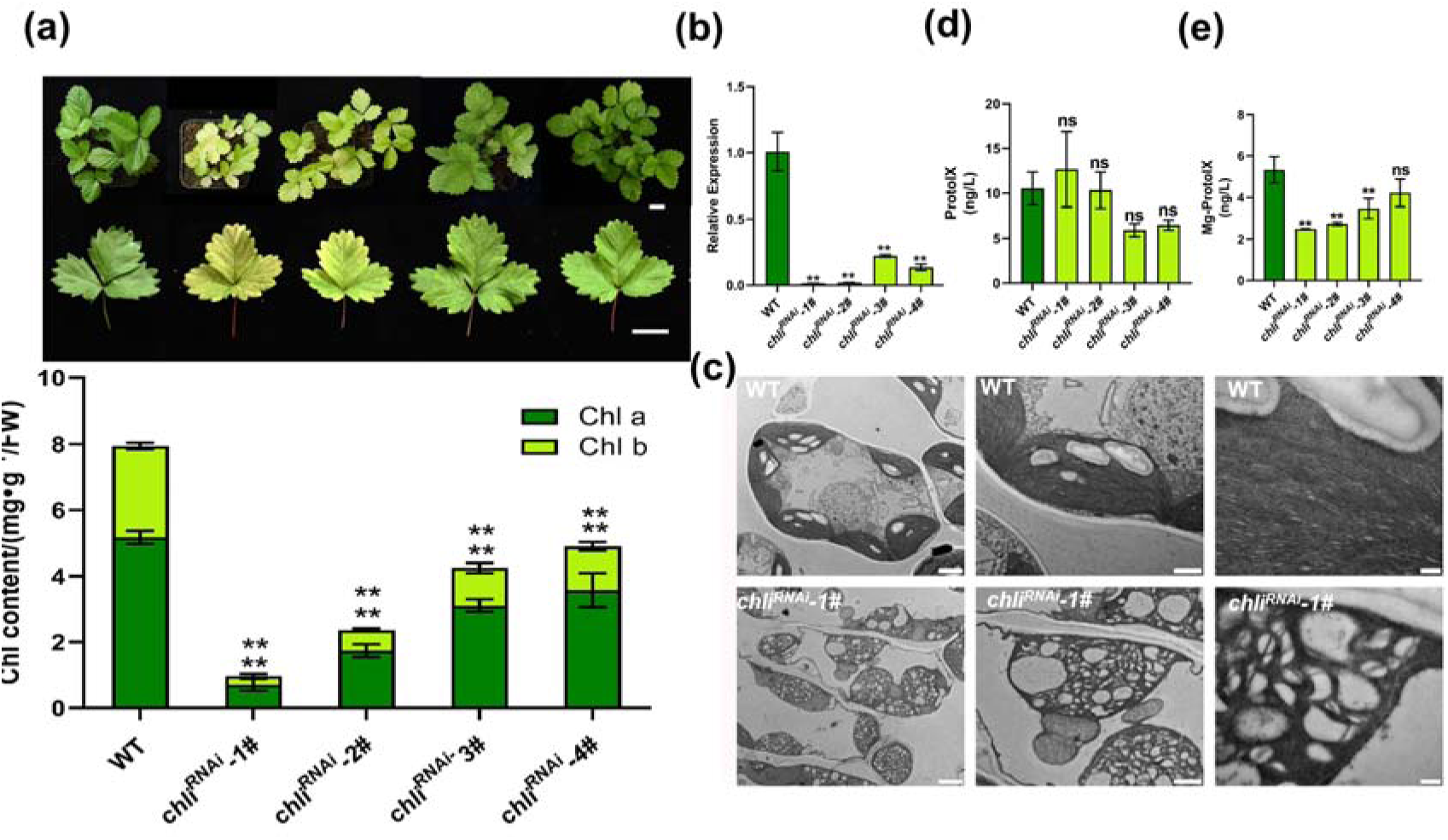
Phenotypes of *chli^RNAi^* in *Fragaria vesca*. (a)Phenotypes of WT and *chli^RNAi^*mutants. Bars,1cm. (b) Relative expression level of *FvCHLI* in *chli^RNAi^*. (c) Proto IX contents of WT and *chli^RNAi^*. (c) Mg-Proto IX contents of WT and *chli^RNAi^*. (d) TEM images shows mesophyll cells and chloroplasts of WT and *chli^RNAi^*, from left to right were mesophyll cells, chloroplasts and magnified chloroplasts. Bars, from left to right, 2μm, 1μm, 200nm, respectively. Data represents the mean±SD of three biological replicates, all data determined by Student’s t-test. **, *p<0.01*; ns, no significant difference.

**Fig. 4.**
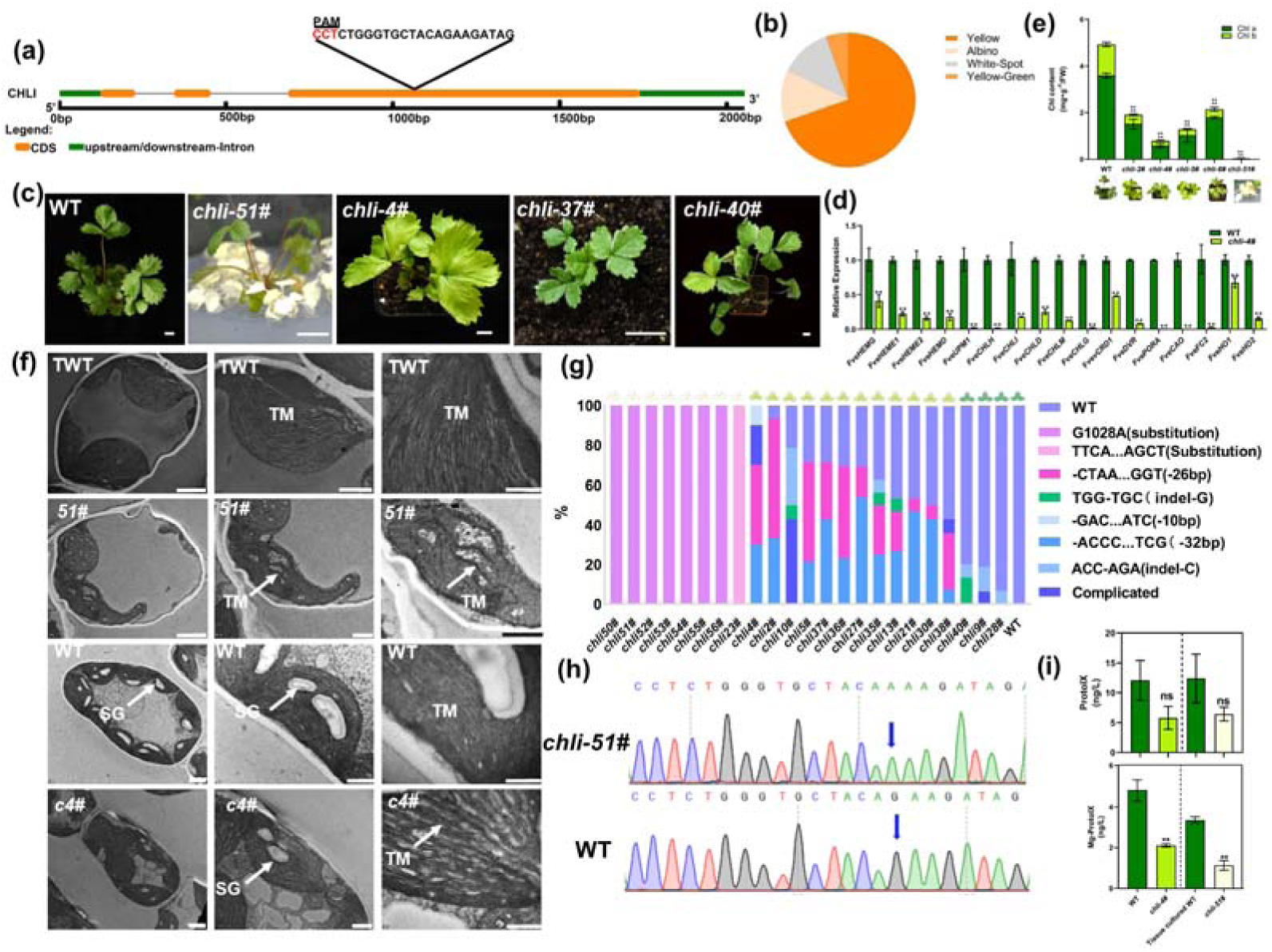
Phenotypes of targeted mutagenesis in *Fragaria vesca*. (a) Schematic map showing the sgRNA target site in *FveChlI*. PAM sequence is marked with red. (b) The percentage of *chli* mutans phenotypes. (c) Phenotypes of WT and *chli* mutants, Bars, 1cm. (d) Reletive expression level of Chl biosynthesis genes. Data analysis using Student’s t-test. **, p<0.01; *, *p<0.05*. (e) Chl contents of *chli* mutants amd WT. Data are mean±SD obtained from triple biological replicates. Data analysis using Student’s t-test. **, *p<0.01*. (f) TEM images shows mesophyll cells and chloroplasts of *chli* mutants and WT, from left to right were mesophyll cells, chloroplasts and magnified chloroplasts. TWT, tissue cultured WT; *51#, chli-51#*; WT, wild type grows on greenhouse; *cs4#, chli-4#*. SG, starch grain, white arrows shows starch grain; TM, thylakoid membrane, white arrows shows thylakoid membrane. Bars, from left to right, 2μm, 1μm, 200nm, respectively. (g) Sequence frequencies in individual plant. Each column represents an individual T_0_ plant where the *FveCHLI* has been targeted. The color codes for sequence mutated is shown on right. Phenotypes corresponding with each T_0_ plant is shown above of each column. (h) Sequencing chromatograms of homozygous mutant. (i) Proto IX and Mg-proto IX contens of *chli* mutant and WT. Data are mean±SD obtained from triple biological replicates. Data analysis using student’s t-test, **, *p<0.01*.

For further analysis of *FpCHLI*, we cloned CDS of *ChlI* from octeploid cultivated strawberry *Fragaria ananassa* Duch cvs.Beijing Taikong 2008, Benihoppe, Ningyu, Sweet Charlie, Akihime and wild diploid strawberry*Fragaria vesca* accessions Hawaii and Ruegen. All PCR products were sequenced. Aligning these sequences suggested that the mutation site was very conserved in these strawberries (Fig. 2c). We also searched NCBI database (https://www.ncbi.nlm.nih.gov/) to collect sequences of other species including *Arabidopsis thalianam, Vitis Vinifera, Solanum Lycopersicum, Malus pumila, Oryza sativa, Thermosynechococcus elongatus and Rhodobacter sphaeroides,* amino acid sequences alignment showed that the 186th amino acid was conserved in all these species (Fig. S5). Hence we preliminary concluded a point mutation of *FpChlI* contributed to the phenotype of *p240*. CHLI is one of three subunits composing Mg-chelatase, which catalysis Proto-IX to Mg-ProtoIX in chlorophyll biosynthesis. Furthermore, Proto IX and Mg-ProtoIX contents have been estimated, not unexpectedly, *p240* possessed less content of Mg-ProtoIX than WT (Fig. 2e), but the same Proto IX as WT (2d), suggesting that *p240* mutant was difficult to synthesis Mg-ProtoIX. These observations provide evidences that mutation made in *FpCHLI* may contribute to the phenotype of *p240*.

### *CHLI* controlled the yellow-green phenotype of *p240*

To assess whether CHLI regulated the yellow-green phenotype of *p240*, we generated *FveCHLI* RNAi plants. We transferred ‘Ruegen’ woodland strawberry via *Agrobacturium*-mediated leaf disk transformation method. The transgenic plants were obtained and verified at RNA level (Fig. 3a and 3b). We marked RNAi plants as *chli^RNAi^*. All *chli^RNAi^* plants indicated a mottled-leaf phenotype, which was highly consistent with RNA level. The transgenic plants which had lower CHLI transcriptional level possessed lower Chl level (both Chla and Chlb) than WT (Fig. 3a). Ultrastructural investigation of chloroplast by TEM suggested the chloroplasts of *chli^RNAi^* were also abnormal (Fig. 3c). In WT, chloroplasts were developed normally, closely stacked thylakoidswere observed. In contrast, *chli^RNAi^* showed no grana stacks in chloroplast. Additionally, numerous vesicle-like structures filled in whole chloroplasts. To investigate whether the Mg-proto IX production was affected, we detected Proto IX and Mg-proto IX contents using a ELISA kits. The result reflects that all mutants presented similar Proto IX level with WT (Fig. 3d). Nevertheless, *chli^RNAi^-1#*,*chli^RNAi^-2#* and *chli^RNAi^*-3# contains significantly lower Mg-Proto IX level than WT, while *chli^RNAi^-4#* has no significant difference with WT (Fig. 3e). These results suggested that CHLI probably plays an important role in phenotype of *p240*.

To further confirm that *ChlI* was responsible for the yellow-green phenotype of *p240*, we also generated *Fvchli* knockout (KO) mutants using CRISPR/Cas9 system. We performed both base editing and genome editing in strawberry. However, we failed to obtain positive plants from base editing. For genome editing, targeted sequences are shown in Fig. 4a. We screened positive T_0_ plants by fluorescence (Fig. S6) and phenotypes. As a consequence, 115 positive T_0_ plants were obtained and 56 (48.7%) were showed obviously different leaf color compared to WT. The phenotypes of 56 T_0_ plants were classified to yellow-green, white-spotted, half-green-half-yellow and albino, respectively. Mutants were treated as chimeras if it bears a mixture of WT (green) and white, which marked white-spotted, and another phenotype half leaf was yellow-green and half leaf was green was marked half-green-half-yellow (Fig. 4c). Finally, among 56 T_0_ plants, we obtained 39 yellow-green (69.6%),7 albinos (12.5%), 7 white-spotted (12.5%) and 3 half-green-half-yellow (5.4%) (Fig. 4b and 4c). Among these mutants, 23 were selected and sequenced. The frequencies and sequence of the various mutations are indicated in Figure 4f. Most of yellow-green mutants generated complicated mutation. WT sequence appeared in 14 mutants while other yellow-green were absence. –CTAA…GGT (indel 26 bp) and –ACCC…TCG (indel 32 bp) mutation events occurred in most yellow-green mutants. White-spotted mutants also performed complicated sequences, but most were WT sequence. For half-green-half-yellow mutants, green tissue generated absolutely WT sequence while yellow showed genotype as yellow-green mutants. Remarkably, six albinos indicated homozygous mutagenesis which possessed a single-base substitution (G to A) in the same mutant position as *p240* (Fig. 4g). We designated these mutants as *chli*. Therefore, we confirm that p240 phenotype was related to mutagenesis in Fp*ChlI*.

Because homozogous mutagenesis showed albinistic leaves and were difficult to survive in the natural environment, we used yellow-green-leaf mutants which had same phenotype as heterozygosis *p240* in our study. For better investigate the functions of CHLI in chlorophyll biogenesis, we determined the transcriptional level of chlorophyll biosynthesis genes, the result suggested that all these genes were down-regulated in *chli4#*(Fig. 4d). Then the pigments of *chli* and WT were measured. Not surprisingly, almost no chlorophyll has been produced for albibino mutant *chli-51#*, while yellow-green mutants *chli-2#, chli-4#, chli-5#, chli-6#* harbored less than half Chla and Chlb of WT. All these mutants were shown significant reduced Chl level compared to WT (Fig. 4e). To investigate whether the defective *FvChlI* affects the chloroplast development in mutants, we compared the ultrastructures of chloroplast in the WT, tissue cultured WT, *chli-4#* and *chli-51#*. We found that both WT and tissue cultured WT had well developed and organized chloroplasts in the mesophyll cells. The closely stacked grana was arranged in the chloroplasts of WT and tissue cultured WT (Fig. 4f). On the contrary, there were almost no stacked grana indicated in chloroplasts of *chli-51#*, and *chli-4#* showed losely stacked grana and significantly decreased starch grain in chloroplasts, which were similar to *p240* (Fig. 4f). In addition to these phenotypes, the concentration of Proto IX and Mg-Proto IX were determined, unsurprisingly, Proto IX level between WT and mutants were showed no significant difference, but all mutants had significantly reduced Mg-ProtoIX (Fig. 4i). These observations demonstrated that the defective *FveCHLI* lead to abnormal chloroplast development and impaired Chlorophyll biosynthesis.

### Missense Mutation of FpCHLI does not affect subcellular localization and the interaction between CHLI and CHLD

The protoplasts of cultivated strawberry ‘TaiKong2008’ were isolated to perform subcellular localization. Both FpCHLI-GFP and Fpchli-GFP were localized with the red autofluorescence of chloroplasts (Fig. S7a). This result indicated that missense mutation of *fpchli* did not affect subcellular localization.

It has been reported that CHLI subunit formed a double-ring complex with CHLD, so whether mutagenesis associated with the interaction between I and D subunit has been verified. The interaction between FpCHLI and FpCHLD was explored by Yeast-two-hybrid system. As show in Fig. S7b, both FpCHLI and Fpchli can bind to FpCHLD, suggesting that though missense mutation was conserved in higher plants, it does not affect the interaction between FpCHLI and FpCHLD.

### Mutation of CHLI result in reduced magnesium chelatase activity and ATPase activity

To further investigate the function of CHLI in strawberry Chl biosynthesis, we predicted the structure of FveCHLI and Fvechli. The protein sequence of FveCHLI was obtained from NCBI database. By changing 186^th^ glutamic acid to lysine, the sequence of *Fvechli* also obtained. Structure prediction performed using Swiss-Model tool and visualized by PyMOL. The predicted structures indicated that mutagenesis amino acid (E186K) decreased the hydrogen bond between 186^th^ and the adjacent amino acid, the hydrogen bonds with VAL189 and GLY191 and disappeared and the LYS186 preferred to bind to HIS230. However, the whole structure was not changed (Fig. 5a and 5b). It was reported that the BchI (CHLI) subunit is responsible for ATP hydrolysis in the catalytic reaction because of its specific structure, ATP binding motifs Walker A (GTGKSTA) and Walker B (LYIDE) (Reid et al., 2004; Walker and Weinstein, et al., 1994). The mutagenesis position of *p240* was close to Walker A and Walker B, so we carried out the ATPase and MgCh activities of mutants and WT, as we expected, both *p240, chli-2#, chli-4#, chli-5#* and *chli^RNAi^-1#* possessed ∼50% of the WT ATPase and MgCh activities under the same condition. The *chli^RNAi^ -3#* and *chli^RNAi^-4#* showed no significant difference with WT in both ATPase and MgCh activities (Fig. 5c-5e). Thus, modified FveCHLI subunits are deficient in ATPase and MgCh, further dampening synthesis of chlorophyll.

**Fig. 5.**
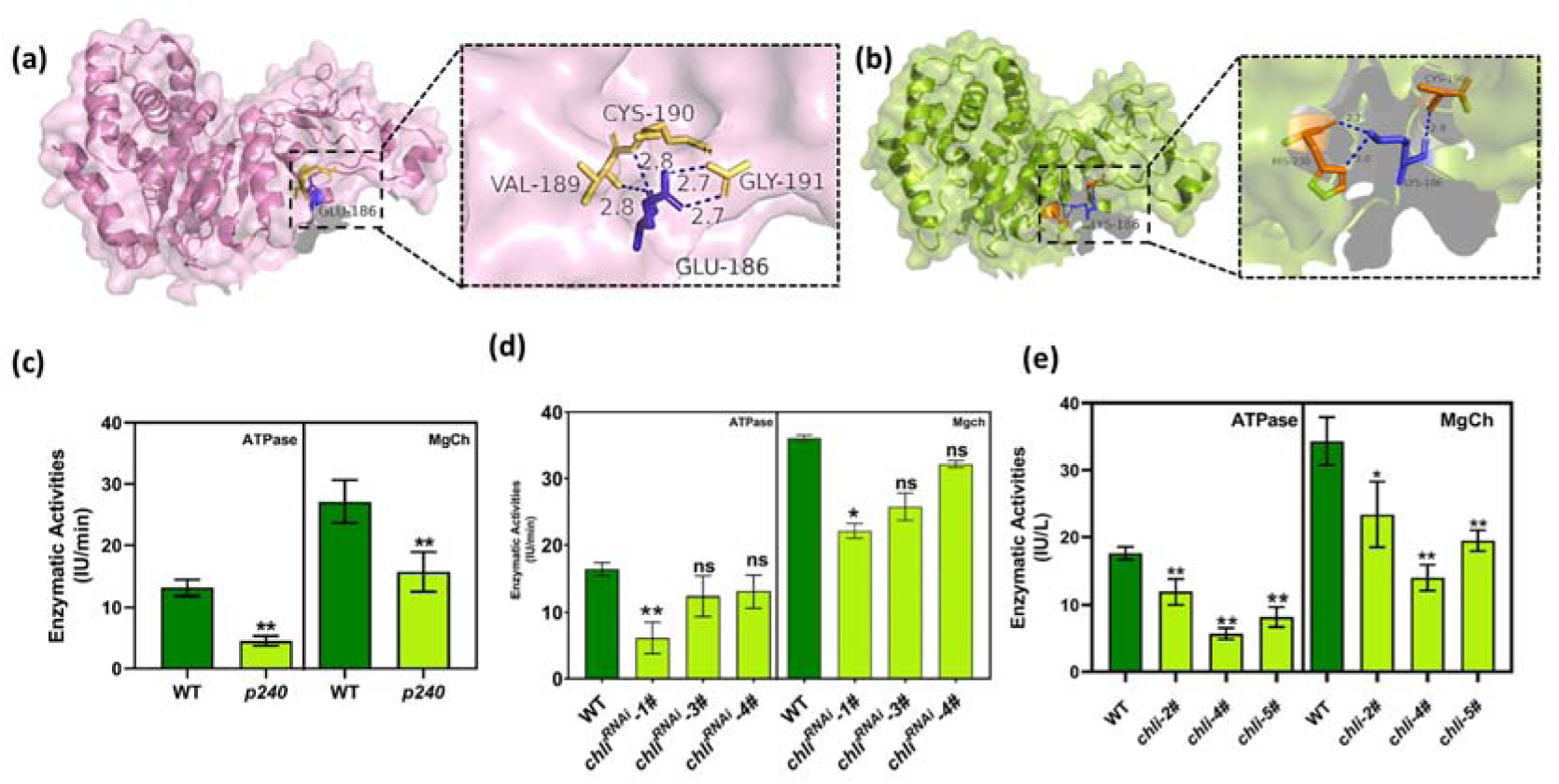
Strcuture prediction of CHLI and enzyme acitivities analysis. (a) Prediction structure of FveCHLI, 186^th^ Glu is marked purple, dotted blue line reprents hydrophobic interaction with other amino acids, number represents magnitude of hydrophobic interaction. (b) Prediction structure of Fvechli, 186^th^ Lys is marked purple, dotted blue line reprents hydrophobic interaction with other amino acids, number represents magnitude of hydrophobic interaction. (c) ATPase and MgCh activities of *p240* and WT, Data are mean±SD obtained from six biological replicates. Data analysis using student’s t-test, **, *p<0.01*. (d) ATPase and MgCh activities of *chli* mutants. Data are mean±SD obtained from triple biological replicates. Data analysis using student’s t-test. **, p<0.01; *, *p<0.05*; ns,no significant difference (e) ATPase and MgCh activities of *chli^RNAi^* mutants. Data are mean±SD obtained from triple biological replicates. Data analysis using student’s t-test. **, *p<0.01*; *, *p<0.05*.

### FveCHLI helps promote flavonoids accumulation in woodland strawberry ‘Ruegen’

The RNA-Seq analysis with leaves from *p240* and WT revealed that 497 of 814 (61.1%) DEGs were related to metabolism, including starch and sucrose metabolism, amino acids metabolism, phenylpropanoid biosynthesis and flavonoid metabolism (Supplementary Fig. S8a). To further characterize whether CHLI influences the metabolite of leaves, we performed nontargeted metabolomics analysis with liquid chromatography–tandem mass spectrometry (LC–MS/MS) using mature leaves in *chli* mutants generated by CRISPR/Cas9(Fig. 6a). Principal component analysis (PCA) revealed that 12 samples were grouped into two independent clusters, indicating that there was a difference between WT and *chli* (Fig. S8b). We totally identified 296 differential metabolites that were annotated in WT and *chli* (Table S4). Clustering analysis indicated that the 296 metabolites comprised 50 flavonoids, 55 lipds, 21 Organoheterocyclic compounds, 36 amino acids and derivatives, 32 organic oxygen compounds, 24 benzenoids, 12 nucleosides_nucleotides_and_analogues, 6 cinnamic acids and derivatives, 5 coumarins and derivatives, 5 stilbenes, 5 alkaloids and derivates, 1 lignans and and 18 other compounds that do not belong to these main classes (Fig.S8d). We screened differential metabolites based on variable importance in projection (VIP)≧1 and a fold change (FC) ≧ 2 or ≤0.5. Totally 250 metabolites were significantly enriched compared to WT, and 98 metabolites accumulated at a lower level in *chli* while 152 were significantly increased in *chli* (Fig. 6b, Table S5). Those metabolites were classified as 42 flavonoids, 46 lipids, 34 amino acids and derivatives, 26 organic oxygen compounds, 25 organoheterocyclic compounds, 17 organic acids and derivatives as well as other compounds (Fig. 6c). KEGG enrichment analysis demonstrated that those metabolites were associated with flavonoid biosynthesis, phenylalanine metabolism and aminoacyl-tRNA biosynthesis (Fig.S8c).

**Fig. 6.**
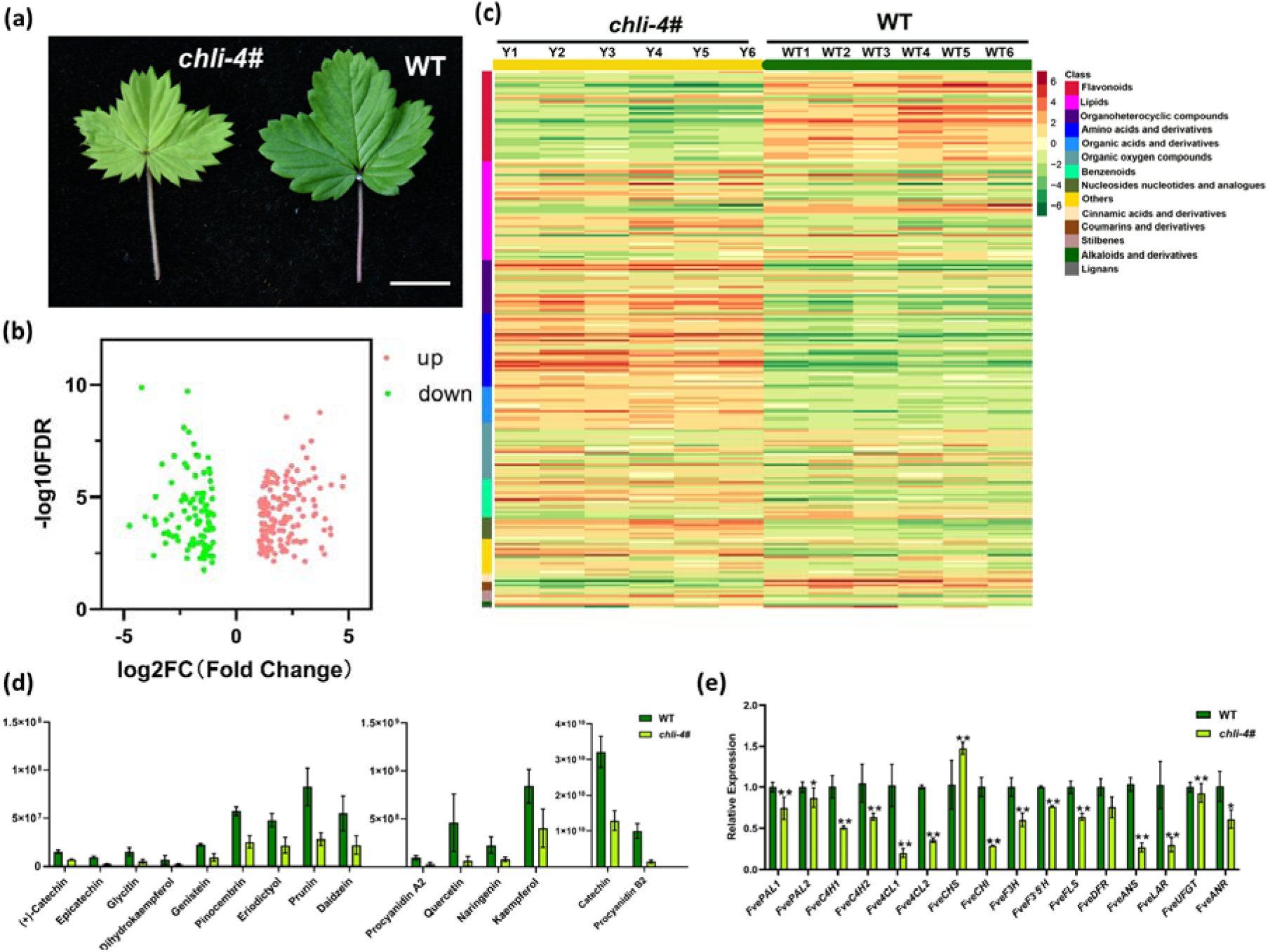
Metabolome analysis of WT and *chli* mutants in mature leaves. (a) leaf phonotype of mature leaves, bar=1cm. (b) The number of metabolites differentially accumulated in *chli* mutant. (c) Clustering heatmap of 251 significantly differential metabolites in *chli* and WT. (d) Relative content of flavonoids in strawberry leaves. Data are mean±SD obtained from six biological replicates. (e) Relative expression level of major genes inflavonoids biosynthesis pathway. Data are mean±SD obtained from triple biological replicates. Data analysis using student’s t-test. **, p<0.01; *, p<0.05

According to the metabolism analysis results, flavonoids were the main differential metabolisms in *chli* and WT, so we focused on flavonoid biosynthesis. Flavonoids were generated from phenylalanine and existed in many subclasses (Fig. S8e). The most abundant flavonoid in wild strawberry was catechin, which belongs to flavanol (Fig. 6d and Table S5). In *chli*, the contents of most (38/42) flavonoids were lower. The major flavonoids, catechins, queroetin, naringenin, and daizein which can scavenge reactive oxygen species directly were all decreased (Fig.6d). In addition, we detected the transcript level of key genes related to flavonoid biosynthesis by qRT-PCR. The results showed that most of these genes were down-regulated in *chli* (Fig. 6e). Thus, CHLI plays a crutial role in maintain flavonoid levels in strawberry leaves.

### *chli* mutants was light-sensitive under low light condition

We noticed *p240* was light sensitive, with yellow leaves under strong sunlight, while pale green leaves under shading condition when cultivated in the field (Fig. S9). To confirm whether heterozygous *chli* has the same phenotype, we treated *chli* with different light density. We obtained light saturation point and light compensation point from light response curve. The *chli* mutants exhibited a lower light compensation point at 350 μmol · m^-2^·s^-1^ while the light compensation point was similar with WT (Table.S7). Since all WT and *chli* mutants were treated with intense light with 300 μmol · m^-2^·s^-1^ and weak light with 50 μmol · m^-2^·s^-1^ for 7 days in a plant growth chamber and (Fig. 7a). We measured the chlorophyll contents using a SPAD-502 meter under different light density. Under intense light with 300 μmol · m^-2^·s^-1^, *chli* mutants accumulated less chlorophyll than WT. However, chlorophyll contents under weak light with 50 μmol · m^-2^·s^-1^ in *chli* mutant were similar to that in WT (Fig. 7b).

**Fig. 7.**
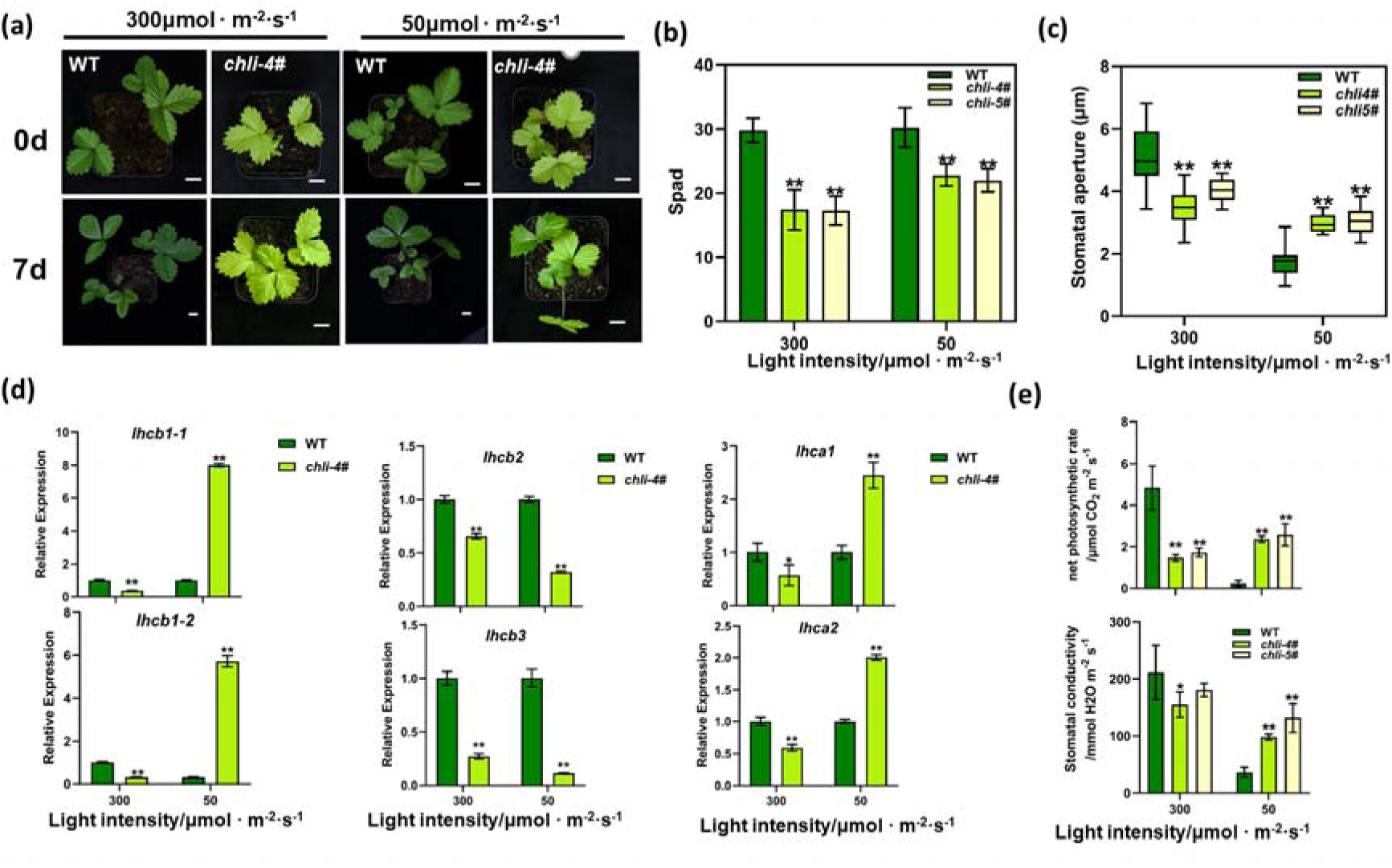
The *chli* mutant exhibits more chlorophyll accumulation under low light condition. (a) The leaf color changes in *chli* under different light indensity for 7 days, bar=1cm. (b) The relative content of chlorophyll in *chli* and WT. Data are mean±SD obtained from ten biological replicates. Data analysis using student’s t-test. **, *p<0.01*. (c) The stomatal apertures under different light indensity. Data are mean±SD obtained from twenty biological replicates. Data analysis using student’s t-test. **, *p<0.01.* (d) Transcriptional level of photosynthetic genes in LHCⅠ and LHCⅡ. Data are mean±SD obtained from three biological replicates. Data analysis using student’s t-test. **, p<0.01; *, p<0.05, ns, no significant difference. (e) Net photosynthetic rate and stomatal conductivity in *chli* and WT. Data are mean±SD obtained from triple biological replicates. Data analysis using student’s t-test. **, p<0.01; *, p<0.05.

We then assessed whether light intensity affect photosynthesis. Net photosynthetic rate and stomatal conductance were determined by CIRAS-3 photosynthesis equipment. We found that photosynthetic capacity in WT was significantly declined while *chli* mutants increased (Fig. 7d). Notably, the stomatal conductance in *chli* was significantly higher than WT under low light (Fig. 7d). We further measured stomatal apertures in the abaxial epidermis microscopically. As we expected, *chli* mutants showed ∼41% wider stomatal apertures than WT plants in shading, but these stomata opened narrower than WT when exposed to light illumination (Fig. 7c and S10). We conclude that *chli* mutant can bear the weak light. We then assessed the expression of several photosynthesis genes related to light harvesting pigment protein complexe(LHCⅠ and LHCⅡ). *Lhcb1* in LHCⅡ, *lhca1* and *lhca2* in LHCⅠ was lower in *chli* mutant under intense light, but higher than WT on weak light. *Lhcb2* and *Lhcb3* in LHCⅡexhibited similar expression pattern, which were reduced both on high and low light. (Fig. 7e). Therefore, we conclude mutated CHLI can improve the low light tolerance of strawberry to some extent.

## Discussion

Based on the chlorophyll-defective leaves, a number of CHLI-associated leaf pigmentation mutants have been reported. Because of diverse mutagenesis position, the phenotype was diverse (Hansson et al., 1999; Huang and Li, 2009; Gao et al., 2016; Dong et al., 2020). In this study, we isolated one yellow-green leaf mutant from ENU mutagenized population of *F. pentaphylla*, which was shown to be caused by a heterozygosis mutation (E186K) of *CHLI* gene. To our knowledge, this chlorophyll-defective mutant is the first chlorophyll biosynthesis mutant in strawberry, and in horticultural crops, only cucumber has been reported to be golden leaf mutant (Gao et al., 2016). According to the phenotype of homozygosis mutants generated from CRISPR/Cas9 mediated gene editing, this mutagenesis position plays an important role in chlorophyll biosynthesis and chloroplast development. Homozygosis mutant displayed albino and lethal, made few chlorophyll and impaired chloroplasts (Fig. 4). However, this mutation neither did effect on subcellular location, nor on interaction between FvCHLI and FvCHLD (Fig. S7). The MgCh and ATPase activities were significantly decreased. In addition to chlorophyll metabolism, FveCHLI affected the accumulation of flavonoids and the response to low light. Metabolome analysis exhibited lower flavonoids level in *chli* mutants, supporting that CHLI helps promote the biosynthesis of flavonoids. Under low light, *chli* mutants were accumulated more chlorophyll than it under high light, revealing that CHLI plays a negative role in light response in strawberry. Overall, our work thus revealed that CHLI also plays an important role in other activities, such as flavonoids metabolism and light response.

The homozygous mutants had an albino phenotype and accumulated few chlorophyll, it cannot survive until cultivated via tissue culture, while heterozygous mutants showed a yellow-green phenotype. These results agreed with Arabidopsis *cs215* mutant (Huang and Li, 2009), barley *Chlorina-125, -157*, and *-161* (Hansson et al., 1999) and maize *Oy1-N1989* (Sawers et al., 2006), which were all albino and lethal but were different to cucumber C582 (Gao et al., 2016) and wheat *ygl1*(Dong et al., 2020) which generated golden leaves and could grow in greenhouse. These diverse phenotypes might be caused by mutated site. Our Glu-186 is only 4 amino acids upstream from the mutations *Chlorina-161* and *Oy1-N1989* mutants, and 6 amino acids downstream from the mutation *cs215*(Fig. S5), suggesting this region was highly conserved. Secondary structure shows this region is an α2 helices and all these mutants in this region resulting an albino phenotype. This region is very important for chlorophyll biogenesis. Previous study showed that the development of thylakoid membranes and chloroplast were related to chlorophyll synthesis (El-Saht, 2000), lower accumulation of chlorophyll in photosynthetic organs resulted abnormal chloroplast development. Our results show that mutation blocks the formation of thylakoid membranes, which corroborate the findings of a great deal of previous work in Zhang (2006).

Previous studies demonstrated that CHLI can interact with CHLD but not CHLH (Zhang et al., 2015), some mutagenesis of CHLI may dampen the interaction between CHLI and CHLD. In our study, though our mutated site is 3 amino acids downstream from rice *ell* mutant, our result is completely opposite, suggesting that FpCHLI also can interact with FpCHLD. However, Mg-chelatase catalysis is a complex process, before CHLI binds to CHLD, they both forms oligomeric ring structures by themselves. ATP is an important molecule for CHLI to a form oligomeric ring structure. Study on crystal structure of *Rhodobacter capsulatus* BchI showed that like other AAA+ protein, BchI forms oligomer in the presence of ATP (Fodje et al., 2001; Lundqvist et al., 2010), then BchI binds to BchD. Therefore, we also predicted the structure of FpCHLI and Fpchli, the results demonstrated that FpCHLI and Fpchli structures were not the same, we presume that FpCHLI can form stable complex with the same binding sites as BchI while Fpchli canno. Hence, we were reasonable to assume that ATPase and MgCh activity might be altered and we confirmed that heterozygous Fpchli decreased ATPase and MgCh (Fig. 5). A similar pattern of results was obtained in barley. Single amino acid substitution of barley in BchI resulted ATPase-deficient BchI, which did not affect forming BchI oligomers, but had no magnesium chelatase activity (Hansson et al., 2002). Based on these studies, though we abandoned to test ATPase and MgCh activity of homozygous mutants because of poor simple quantities of homozygous Fpchli, we can hypothesis that homozygous Fpchli displays few ATPase and MgCh activities.

Flavonoids are a group of polyphenol compounds that widely distributed in plant vacuole chloroplast and nucleus. These natural substances play important roles in response to many biotic and abiotic factors. Queroetin, naringenin, catechins and daidzein were main flavonoids that play a role in stabilization and scavenging of ROS directly (Shen et al., 2022). These substances were accumulated in our wild type strawberry and reduced in KO mutant, suggesting that yellow-leaf mutants produced lower flavonoids level. The same result was found in two albino tea cultivar ‘Anjibaicha’ and ‘Huangjinya’, which all showed reduced flavonoids especially lower catechins. However, the expression of major flavonoids biosynthesis genes were up-regulated in these tea cultivars (Song et al., 2017). In Our study, flavonoids biosynthesis genes were down-regulated except chalcone synthase (*CHS*) in *chli* mutant. Wei (2011) reported that the rise of chlorophyll a content in tea was consistent with the increase of (-)-epicatechin, (-)-epigallocatechin and the decline of (+)-catechin, suggesting that chlorophyll plays a vital role in regulation of catechin. Our study also provided an evidence that chlorophyll level has a correlation with flavonoids, and low chlorophyll level was related to low catechin. However, the detailed mechanism by which *FveCHLI* influences flavonoids accumulation in strawberry leaves awaits further research.

We found heterozygous *p240* mutant is light-sensitive, with a phenotype of yellow leaves in sunlight but green in shade. Subsequently, our heterozygous KO mutants showed the same phenotype as *p240*. The same reports were found in tea plants, as albino-leaf plants ‘Huangjinya’ and ‘Anjibaicha’ showed a yellow-leaf phenotype under strong light but green leaf under weak light (Song et al., 2017; Wei et al., 2011). Further study showed that shading might increase the expression of *CsPORL-2* (encoding protochlorophyllide oxidoreductase) by inhibiting the expression of CsHY5, resulting an increase of chlorophyll in tea leaves (Chen et al., 2021). However, how FveCHLI correlation with low light and accumulation of chlorophyll should be investigated further.

In summary, in addition to common role in chlorophyll biosynthesis, our results reveal additional functions of FveCHLI at leaves metabolism and light response. Overall, we show not only that FveCHLI is important for chlorophyll metabolism, but also the regulation of FveCHLI is a promising strategy to breed low light tolerant varieties.

## Materials and Methods

### Plant Material and Growth Conditions

The white-fruited *Fragaria pentaphylla* assession ‘Pao’er’ and red-fruited *Fragaria vesca* assession ‘Ruegen’ were both used as wild types in this study. The *p240* mutant was obtained by ENU mutagenesis of *F. pentaphylla* seeds. Mutagenesis was performed by Luo et al. (2018). Briefly, seeds were soaked in 0.4% ENU for 7h at room temperature with gentle shaking, then sterilized and cultured on 1×Murashige and Skoog (M519, Phytotech) medium. After one months’ cultivation, seedlings were all grown in a plant growth room for screening.

All strawberries were cultivated for one months in a growth room under a 16h photoperiod at 25℃ temperature and a light intensity of 300 μmol m^-2^s^-1^, then transplanted in a greenhouse at Northwest A&F University, China, Yangling.

### Pigments, Chl Precursor and enzyme activity Analysis

Chlorophyll was extracted from fresh strawberry leaves and determined as previously described by Lichtenthaler (1987). Chlorophylls and carotenoids were extracted using 80% acetone and measured by an ultraviolet spectrophotometer (UV-3802S, Shanghai, China). The following formulas were used to quantify the pigments:

Concentration of chlorophyll a (C_a_)=(12.25A_663_-2.79A_647_)×v/w

Concentration of chlorophyll b (C_b_)=(21.50A_647_-5.10A_663_)×v/w

Concentration of carotenoids (C_x+c_)=((1000A_470_-1.82C_a_-85.02C_b_)/198)×v/w

(V and W represents the volume of acetone and the fresh weight of leaf samples, respectively).

The precursor Proto IX and Mg-Proto IX were quantified using Plant ProtoIX/Mg-Proto IX ELISA Kit (HS354-Pt, Hengyuan, Shanghai, China). ATPase and magnesium chelatase (MgCh) activities were performed using Plant ATPase (HB328X-Pt, Hengyuan, Shanghai, China) and MgCh ELISA Kit (HB144X-Pt, Hengyuan, Shanghai, China).

Chl precursor and enzyme activities were shared same leaf pretreatment. Approximately 0.5 g fresh strawberry leaves were homogenized in PBS (pH7.2), about 1 g sample added 9 ml PBS, then centrifuged at 3000 g for 30 min at 4℃. The supernatants were collected in a 10 ml tube. Finally, the supernatants were treated using manufacture’s instruction. After treatment, measure the absorbance (OD) of each sample at 450 nm using a microplate reader (Infinite M200pro, Tecan, Switzerland).

### Transmission Electron Microscopy

Wild type and mutants leaf sections were fixed in 2.5% glutaraldehyde at 4°C overnight, and further fixed in 1% osmic acid, dehydrated with 30%, 50%, 70%, 80%, 90% and 100% ethanol for 10 min each, then stained with 2% uranium dioxy acetate and Lead citrate trihydrate, and photographed under a transmission electron microscope (TECNAI G2 SPIRIT BIO, FEI, USA). The number of starch grain was calculated for 10 cells of each sample.

### Protoplast isolation and chloroplast calculation

Mesophyll protoplasts of wild type and *p240* mutant were isolated as described by Gou (2019), and observed using an Olympus BX-51 fluorescence microscope (BX51, Olympus, Japan). Each sample took 20 protoplasts for chloroplast accounting.

### RNA-seq analysis

The transcriptome sequencing was carried out by Beijing, Novogen Com., Ltd (https://cn.novogene.com/). Total RNA was extracted from the leaf samples of WT (*F. pentaphylla*) and *p240* mutant, using the Plant Total RNA Extraction Kit (OMEGA, Bienne, Switzerland). Each sample included three biological replicates. The RNA quality was assessed using both the NanoPhotometer ^®^ spectrophotometer (IMPLEN, CA, USA) and the RNA Nano 6000 Assay Kit of the Bioanalyzer 2100 system (Agilent Technologies, CA, USA). Sequencing libraries were constructed using NEB Next^®^ UltraTM RNA Library Prep Kit for Illumina (NEB, Ipswich, MA, USA) following manufacturer’s recommendations and index codes were added to attribute sequences to each sample. After cluster generation, library preparations were sequenced on the Illumina Novaseq platform to generate 150 bp paired-end reads. Clean reads were mapped to the reference genome *Fragaria vesca* (Hirakawa et al., 2014) using Hisat2 tools. Genes with adjusted P-value <0.05 found by DESeq2 were considered as differentially expressed. SNP calling was performed using GATK2 (v3.7) software. The vcf files were filtered with GATK standard filtering method and other parameters (cluster:3; WindowSize:35; QD < 2.0 o; FS > 30.0; DP < 10), and the Variablesite was annotated with SnpEff software.

### Plasmid construction

Genomic DNA or cDNA used for sequence amplification were obtained from young leaves of ‘Ruegen’. For CRISPR/Cas9, sgRNA was designed containing the mutagenesis region aligned to *p240* (Table S1). The CRISPR/Cas9 vector used for transformation was modified from pKSE401 (Xing et al., 2014). The modification method was performed as described by Tang (2018) with minor differentiation. The 35S-sGFP-terminator cassette was inserted into the *EcoR*Ⅰ site of pKSE401 by the Gibson assembly method (Gibson et al., 2009). The plant expression vector was constructed as described previously (Xing et al., 2014). Plasmid was transformed into *Agrobacterium tumefaciens* GV3101 for genetic transformation.

For subcellular location analysis, the full-length coding sequence of FpCHLI was amplified from F. pentaphylla. The PCR product was digested and fused in pBI221 vector as described by Wei (2016). The Fpchli was obtained by over-lapping PCR. Briefly, two rounds PCR were conducted. In the first round, PCR product was amplified from FpCHLI-pBI221 using specific primer containing mutated site. In the second round, the PCR fragment was reassembled by using one step cloning kit (vazyme, C112). Both PCR products were digested and fused in pBI221 vector as described by Wei (2016).

All primers used are listed in supplemented Table 1.

### Strawberry stable transformation

*Agrobacterium*-mediated transformation was performed as previously described with minor modification (Feng et al., 2019). Briefly, leaf strips of ‘Ruegen’ were cultured on preincubation medium (MS, 2% sucrose,2 mg·L^-1^thidiazuron (TDZ), 0.8 mg·L^-1^ indole-3-butyric acid (IBA), 0.8% agar) for one week in the dark; Socked in the *Agrobacterium tumefaciens* GV3101 carrying each construct (OD 0.4-0.6) for 10-15min with gentle shaking. The tissue was then placed to preincubation medium to co-cultivation for 3 d in the dark. After washing with sterile water three times, the materials were kept on delayed-screening medium (preincubation medium with 200 mg·L^-1^ timentin and 300 mg·L^-1^ cefuroxime) for 2 weeks in the dark, and transferred to screening medium (delayed-screening medium with 15mg·L^-1^kanamycin) under light condition until adventitious buds developed. Positive adventitious buds were selected by GFP fluorescence using a fluorescence dissecting microscope (MZ10F, Leica, Germany) and placed to rooting medium (MS with 3% sucrose, 0.7% agar, 200 mg·L^−1^timentin and 300 mg·L^-1^ cefuroxime). Rooted plantlets were transplanted into pots for further analysis.

### Detection of transgenic-plant

T_0_ transgenic plants were identified by the presence of GFP fluorescence and PCR amplification. Genomic DNA of T_0_ plants was extracted using a CTAB-based method (Allen et al., 2006). The DNA fragments were amplified using primers U6-26-P and U6-26-T (Table S1) to test the sgRNA regions. The target site mutation was identified by amplifying the genomic DNA fragments of T_0_ plants using primers gCHLI-F and gCHLI-R (Table S1). The PCR products were purified using Gel Extraction Kit and sequenced at TSINGKE, China (Xi’an). The results were analysed by DSDecode (http://dsdecode.scgene.com/home/) (Liu et al., 2015) to decode sequence and detect mutations. For complicated results which cannot decode, we transferred purified PCR products to pMD19-T Vector (D102a, Tarkara) and 15 colonies for each plant were sequenced.

### Subcellular Localization assay

To examine subcellular localization, transient expression of *Arabidopsis* mesophyll protoplasts was carried out as described by Wei (2016). The GFP fluorescence was detected by laser scanning confocal microscope (TCS SP8 SR, Leica, Germany).

### Yeast Two-Hybrid Assays

A yeast two-hybrid assay was performed according to the manufacturer’s instruction (PG1031, pytbio). The full-length CDS of *FpGHLI* and *fpchli* were first introduced into pBI221, then cloned to pGBDT7 using primer 221 to pGBDT7. CDS of FpCHLD and FpCHLH were cloned into pGADT7 by the same method. All plasmids were co-transformed into yeast strain AH109. Positive clones were selected on SD/-LeuTrp, and then on transferred onto SD/-Leu-Trp-His-Ade + 40mg/L x-α-gal plates. The primers used are listed in Supplementary Table 1.

### qRT-PCR analysis

Total RNA was extracted from the leaf samples of WT (*F. vesca* assecion ‘Ruegen’), *chli* and *chli^RNAi^* mutants, using the Plant Total RNA Extraction Kit (R6827-01, OMEGA, Bienne, Switzerland). Reverse transcription for first-strand cDNA was performed with HiScript® II Q RT SuperMix (Vazyme Biotech Co., Nanjing, China). *FveCHP1* was used as the reference gene (Clancy et al., 2013). qRT-PCR reactions were conducted with Taq Pro Universal SYBR qPCR Master Mix (Vazyme Biotech Co., Q712-02) using Applied Biosystems StepOne TM real-time quantitative PCR system (Applied Biosystems, USA). The expression of each gene was normalized against with WT. The primers were listed in Supplemented table 1.

### Protein structure prediction

The FASTA sequence of FveCHLI was downloaded from NCBI database. The Fvechli sequence was obtained by changing the 186^th^ glutamic acid to lysine. Both sequences were used for structure prediction. We performed protein folding trajectories using Swiss-Model (https://swissmodel.expasy.org/). The trajectories were visualized via PyMOL (Yuan et al., 2016).

### Metabolome analysis

Six biological repeats were used for metabolome analysis which were performed by Beijing, Novogen Com., Ltd (https://cn.novogene.com/). Mature leaves (100mg) from *chli-4#* and WT were grinded with liquid nitrogen and the homogenate was resuspended with prechilled 80% methanol by well swirl, respectively. After a 5 min incubation on ice, these samples were centrifuged at 15,000g, 4 ℃ for 20 min. Dilute the supernatant to a final concentration containing 53% methanol by LC-MS grade water. Transfer the samples to new Eppendorf tubes and centrifuge at 15,000 g, 4 °C for 20 min. Finally, the supernatant was collected for LC-MS/MS system analysis. The processed samples were analyzed by using a Vanquish UHPLC system (ThermoFisher, Germany) coupled with an Orbitrap Q Exactive^TM^HF-X mass spectrometer (Thermo Fisher, Germany) in Novogene Co., Ltd. (Beijing, China). Samples were injected onto a Hypesil Gold column (100×2.1 mm, 1.9μm) using a 17-min linear gradient at a flow rate of 0.2mL/min. Both positive and negative polarity mode were analyzed. The positive polarity eluents were eluent A (0.1% FA in Water) and eluent B (methanol), and the negative polarity eluents were eluent A (5 mM ammonium acetate, pH 9.0) and eluent B(Methanol). The solvent gradient was set to: 2% B, 1.5 min; 2-100% B, 3 min; 100% B, 10 min; 100-2% B, 10.1 min; 2% B, 12 min. Q Exactive^TM^ HF-X mass spectrometer was operated in positive/negative polarity mode with spray voltage of 3.5 kV, capillary temperature of 320°C, sheath gas flow rate of 35 psi, aux gas flow rate of 10 L/min, S-lens RF level of 60 and aux gas heater temperature of 350°C.

The raw data files generated by UHPLC-MS/MS were processed using the Compound Discoverer 3.1 (CD3.1, ThermoFisher) for peak alignment, peak picking, and quantitation for each metabolite. The normalized data was used to predict the molecular formulas based on additional ions, molecular ion peaks and fragment ions. Peaks were then matched against mzCloud (https://www.mzcloud.org/) mzVault and MassList database for accurate qualitative and relative quantitative results. Statistical analyses were performed using the statistical software R (R version R-3.4.3), Python (Python 2.7.6 version) and CentOS (CentOS release 6.6), When data were not normally distributed, try using the area normalization method for normal transformations.

Principal components analysis (PCA) and Partial least squares discriminant analysis (PLS-DA) were performed on metaX, a flexible and comprehensive metabolomics data processing software. The hierarchical clustering analysis was performed using the heat map method. The data were normalized with z-scores of regions of differential metabolite intensity and plotted with the R language Pheatmap package. The significantly differential metabolites between groups were calculated as VIP > 1 and P-value< 0.05 and fold change≥2 or FC≤0.5. These metabolites were annotated through the KEGG database (https://www.genome.jp/kegg/pathway.html), the HMDB database (https://hmdb.ca/ metabolites) and LIPIDMaps database (http://www.lipidmaps.org/). Metabolic pathway enrichment was considered as statistically significant when its P-value < 0.05.

### Light intensity treatment

The light response curve measurement was performed using LI6800 Portable Photosynthesis System ((LiCoR, Lincoln, NE, USA)) using functional leaves of WT and chli mutants. Measurements were conducted following a series of decreasing PPFDs from 1600, 1400, 1200, 1000, 800, 600, 400, 300, 100, 80, 50 and 0 μmol m^-2^s^-1^. The constant CO_2_ concentration were 400 μmol m-^2^s^-1^μmol. The light response curve was done by Phtotosynthesis software. Three replicates were taken for each response curve.

For different light intensity treatment, all mutants and WT were cultivated in a growth chamber with light intensity of 300 μmol m^-2^s^-1^. For weak light treatment, mutants and WT were shaded by a single layer shade net which reduced light intensity to 50 μmol m^-2^s^-1^. After 7 days, chlorophyll contents were measured by a SPAD-502 meter. The SPAD-502 meter is a portable device which is widely used for chlorophyll measurement rapidly, accurately and non-destructively. The relative SPAD meter values that are proportional to the chlorophyll content in the leaf (Ling et al., 2011). Photosynthetic parameters were measured using the CIRAS-3 system (PPSYSTEMS, USA), and parameters were set and calculated with manufacturer’s instruction and software. Stomatal aperture measurement was measured according to a previous method. The transparent tape was applied to the upper epidermis of strawberry leaves, and the mesophyll was carefully scraped off with a clean surgical blade. Then the lower epidermis was stained with potassium iodide for 30s, then cleaned with sterile water, and re-attached to glass slides. The stomata were observed with BX51(Olympus, Japan) and photographed. Stomatal apertures were measured by ImageJ software. Twenty stomata were measured for each treatment. Meanwhile, leaves at same position were harvested and frozed in -80℃ ultra low temperature refrigerator for qRT-PCR analysis.

### Statistical analyses

Statistical analyses were conducted by SPSS v26.0 (IBM, USA). Student’s t test (\**P*<0.05, \*\**P*< 0.01), was used to perform pairwise comparisons.

## Author and Contributors

JYF and YW conceived the study; YM, JS, DW XL, and FW performed the experiments; YM wrote the manuscript; and JYF, YW, CG, LQ and YM interpreted the experimental data and revised the manuscript. All of the authors read and approved the final manuscript.

## Acknowledgements

We would like to thank the anonymous reviewers for comments on the manuscript. The present study was financially supported by grants from the project Scientific and technological innovation and achievement transformation project of Northwest A&F University experimental demonstration station (202102) to J.Y. F., and Major projects of Agricultural Cooperative Innovation and Extension Alliance in Shaanxi Province support (LMZD202102) to Y.Q. W.

